# Assessing the Stability of Molecular Glues with Weighted Ensemble Simulations

**DOI:** 10.64898/2026.01.23.701371

**Authors:** Seref Berk Atik, Alex Dickson

**Affiliations:** Department of Computational Mathematics, Science and Engineering, Michigan State University, East Lansing, Michigan 48824, USA; Department of Biochemistry and Molecular Biology, Michigan State University, East Lansing, Michigan 48824, USA

## Abstract

Targeted protein degradation is an emerging approach that utilizes cellular degradation pathways to inhibit a target protein. Small molecules such as molecular glues or PROTACs can be used to mediate the formation of a ternary complex with an E3 ligase and the target protein, which can dramatically enhance the degradation process. This approach is promising for cancer therapy, where degradation of oncogenic proteins can lead to cancer cell toxicity. To design new molecular glues, it is important to develop methods that predict how well a given molecule stabilizes a protein-protein interaction. However, conventional molecular dynamics simulations face challenges in capturing the long-timescale binding and unbinding events that would be used to evaluate this stabilization. In this study, we developed a strategy that allows us to evaluate the stability of protein-protein interactions in the presence of a glue molecule using weighted ensemble simulations in combination with weakened protein-protein interactions. Using this strategy, we generated unbinding trajectories of the DCAF15-RBM39 system with small molecules E7820, Indisulam, and several other Indisulam analogs. We were able to observe distinctly different behaviors between systems with different glues, which was in agreement with their reported EC_50_ values. We believe this approach could aid drug discovery efforts by expanding the set of druggable targets and improving the success rate of molecular glue development.

## Introduction

Inhibiting the function of disease-associated proteins has long been a crucial strategy in therapeutic development. Traditionally, this has been achieved using small-molecule inhibitors that bind to well-defined pockets on target proteins, often active or allosteric sites. However, the effectiveness of these compounds is limited by their requirement for persistent occupancy and the necessity of a suitable binding site, rendering many proteins “undruggable” by conventional means. Targeted protein degradation (TPD) offers an alternative approach by harnessing the cell’s ubiquitin–proteasome system (UPS) to eliminate, rather than inhibit target proteins [57, 44, 2, 56, 49]. UPS can be broadly divided into four main components: the E1 ubiquitin-activating enzyme, the E2 ubiquitin-conjugating enzyme, the E3 ubiquitin ligase and the 26S proteasome [4]. The E3 ligase allows substrate specificity by binding to the target protein, positioning it for the transfer of ubiquitin from E2 [31]. Once the substrate is ubiquitinated, it is recognized and degraded by the proteasome. Molecular glues [45, 35] and proteolysis-targeting chimeras (PROTACs) [42, 12], both referred to as degraders, can exploit this pathway by promoting the formation of a ternary complex between an E3 ligase and a protein of interest (POI). This interaction promotes the ubiquitination and subsequent degradation of the target protein by the proteasome [11]. After the target protein is ubiquitinated, the degrader molecule is released from the complex and can continue to engage in the formation of other ternary complexes, thus enhancing its overall potency [43]. Without the need for conventional binding pockets, the mechanism of action of degraders allows for the targeting of previously undruggable proteins [9].

Among the various TPD strategies, molecular glues gained attention due to their pharmacological advantages [55]. In contrast to PROTACs – which engage the POI and E3 ligase with separate moieties that are connected by a flexible linker – molecular glues are small molecules that induce proximity by filling gaps in the protein-protein interface. This smaller size allows them to have more favorable pharmaceutical properties in terms of increased oral bio-availability and improved cellular permeability relative to PROTACs [55, 35]. The ability to bind tightly to both ligase and a POI also makes PROTACs susceptible to the hook effect [19, 58], where POI-ligase pairs are unable to coordinate as they are both bound to separate PROTACs.

The discovery of molecular glues has traditionally relied on observation-based approaches, with several early examples identified incidentally rather than through rational design [46]. More recently, efforts have shifted toward systematic exploration, where analogs of known glue molecules are synthesized and evaluated to identify new candidates [43]. High-throughput methodologies, particularly microarray-based screening platforms, have enabled the screening of large chemical libraries to uncover compounds capable of promoting targeted degradation. These experimental strategies are often integrated with biological validation assays and medicinal chemistry optimization to refine initial hits for potency and specificity [47, 40].

While these workflows have improved discovery rates, the identification, characterization, and optimization of molecular glue candidates can greatly benefit from a detailed understanding of the ternary interface and its conformational dynamics. It is now possible to acquire such information through structure-based methods such as docking [6, 24], molecular dynamics (MD) simulations [13], and free-energy calculations. In combination with deep-learning based tools [51, 37], physics-based methods enable the identification of key interface residues and the sampling of the relevant conformation space [17, 16, 1, 33, 39]. In recent research on molecular glues, MD simulations have been used to investigate conformational dynamics and ternary complex stability [54, 51, 5]. However, conventional MD simulations are often limited in their ability to capture rare events such as ternary complex dissociation, which may occur on timescales beyond practical simulation windows [52]. To overcome this limitation, enhanced sampling strategies have been developed to accelerate the exploration of conformation space while preserving thermodynamic accuracy. Among these, the weighted ensemble (WE) method [28] offers a framework to efficiently sample rare events by distributing computational effort over parallel trajectories. This approach enables the observation of rare but functionally relevant events without biasing the system dynamics [53]. In this study, WE simulations were performed using the Wepy toolkit, which is an open-source Python framework for running and analyzing WE simulations [38]. The Resampling of Ensembles by Variation Optimization (REVO) algorithm was used to enhance trajectory diversity and improve sampling efficiency [18]. This simulation approach is applied to investigate the structural stability of the ternary complex systems.

To evaluate the application of this WE–based strategy, we selected the DCAF15–RBM39 system, a well-established model in molecular glue research [30]. As for molecular glues, Aryl sulfonamide compounds, including E7820 and Indisulam have been shown to promote selective degradation by inducing ternary complex formation between DCAF15 and RBM39 [26, 3, 10, 20, 22, 7]. Structural studies have demonstrated that these compounds engage at a shallow surface pocket on DCAF15 facilitating the recruitment of RBM39 through its RRM2 domain [20, 10, 22]. This interaction is not driven by intrinsic affinity between the ligand and E3-ligase alone, but is stabilized by a network of complementary protein–protein, protein-glue and water-mediated interactions across the ternary interface [20, 10, 22]. In this study, we investigate a series of six aryl sulfonamide molecular glues, including E7820, Indisulam, and structurally related analogs reported by Bussìere et al [10]. These compounds span a range of biochemical potencies, from 0.74 µM to undetectable [10], providing a strong evaluation of our enhanced sampling simulation pipeline.

## Methods

### Molecular Glues

We investigate a series of six molecular glues designed to promote ternary complex formation between DCAF15 and RBM39. We selected E7820, a potent degrader, along with Indisulam and several of its analogs characterized in a previous study by Bussìere et al [41, 10, 22]. To ensure that our method was evaluated across a spectrum of activity levels, we included compounds 4, 5, 7, and 10 from the aforementioned study, hereafter referred to as C4, C5, C7, and C10, respectively (Figure 1). An apo system, without any glue molecule, was also included as a control.

**Figure 1:**
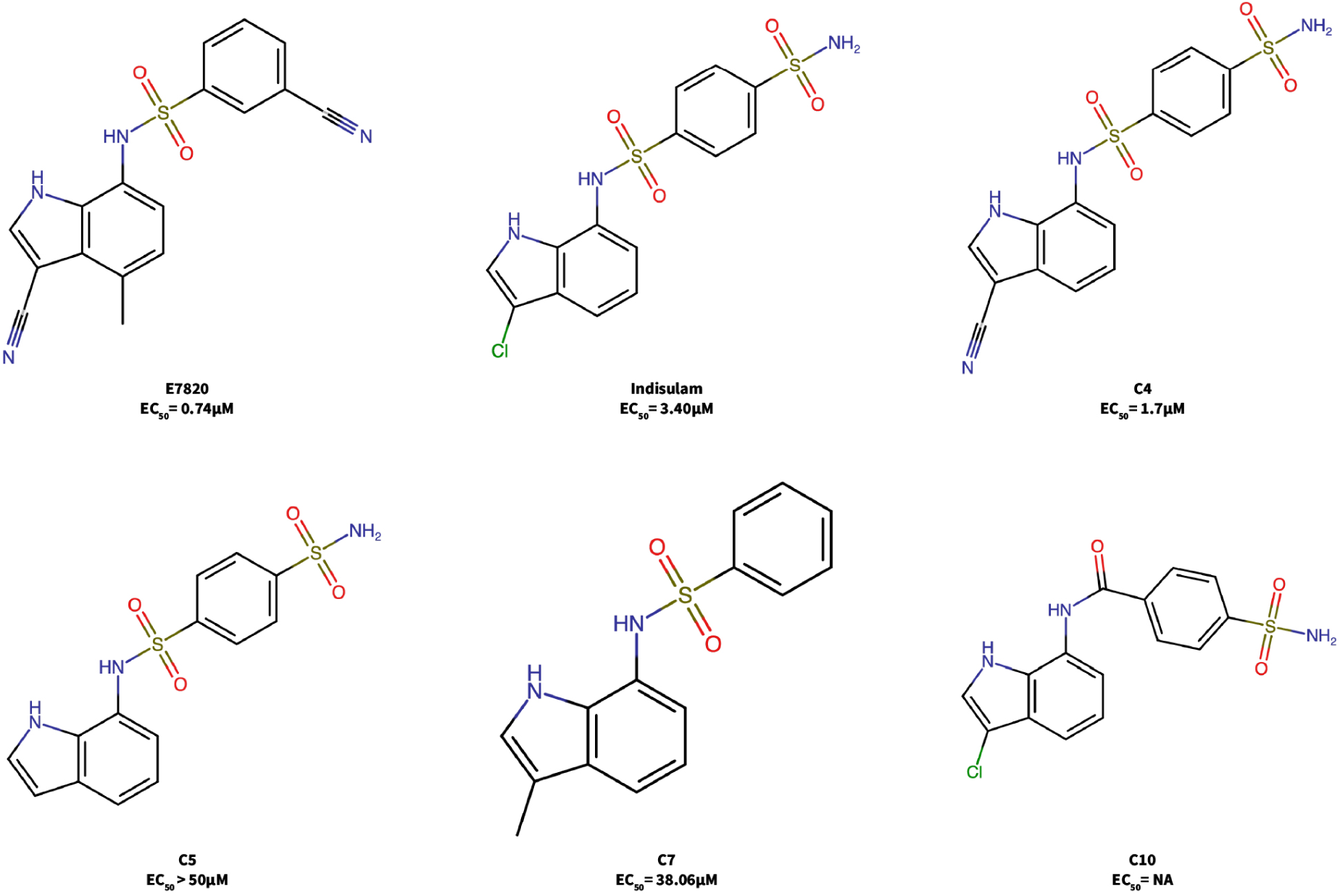
Chemical structures of the molecular glues with their corresponding EC_50_ values: E7820, Indisulam, C4, C5, C7, and C10 [10, 22].

### System Preparation

The starting structure for the ternary complex of DCAF15, RBM39, and the molecular glue E7820 was obtained from the Protein Data Bank (PDB ID: 6PAI), which includes all three components in a crystallized conformation [20]. This structure served as the template for generating alternative systems by replacing E7820 with other glue candidates (e.g., Indisulam, C4, C5, C7, C10) using the CHARMM-GUI Ligand Reader & Modeler module [34]. This module preserves the shared heavy-atom positions of the original molecular glue and enables construction of a modified glue molecule directly within the binding pocket. Each ternary complex was solvated in a TIP3P water box using the CHARMM-GUI Solution Builder [32, 8] and neutralized with 0.15 M NaCl. Force field parameters were assigned using the CHARMM36m force field [27] for proteins and the CHARMM General Force Field (CGenFF) [50] for the small-molecule glues, generated by Ligand Reader & Modeler [36]. To improve simulation efficiency, Hydrogen Mass Repartitioning (HMR) [23] was enabled, allowing a time step of 0.004 picoseconds during MD simulations.

### Molecular Dynamics Simulations

All systems were first energy minimized to relax particle positions. Energy minimization was performed with a convergence tolerance of 100 kJ/mol/nm over a maximum of 5000 steps. During minimization, positional restraints are applied to the protein with force constants of 400 kJ/mol/nm^2^ for backbone atoms and 40 kJ/mol/nm^2^ for side chains. Following minimization, initial velocities were assigned according to a Maxwell-Boltzmann distribution, and the system was gradually heated to 303.15K over 125 ps with a timestep of 1 fs in the canonical (NVT) ensemble. All dynamics were run using OpenMM v8.0 [21] with the Langevin Integrator and a friction coefficient of 10 ps*^−^*^1^.

After the equilibration, the non-bonded interaction strengths, specifically electrostatic and Van der Waals forces, between DCAF15 and RBM39 were scaled down by adding an energy term (E_rep_), defined as follows:

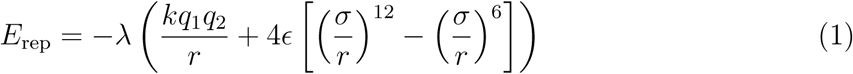

This is simply the negative of the standard non-bonded interactions computed between the proteins, scaled down by a factor λ ∈ [0, 1]. The energy term was defined using an OpenMM CustomNonbondedForce that was only applied to inter-protein atom pairs; this did not affect intramolecular protein interactions. Particle parameters (charge and type) and tabulated functions were copied directly from the original CustomNonbondedForce to preserve consistency, and exclusions were maintained. A range of scaling factors were systematically tested to identify a balance between preserving structural stability and allowing sufficient sampling of protein unbinding pathways. The optimal factor enabled the exploration of glue-dependent stability while avoiding complete destabilization of the complex. Introducing this scaling allowed us to accelerate the sampling of dissociation events within the ternary complex, thereby enhancing the efficiency of subsequent WE simulations.

### Weighted Ensemble Simulations

Following equilibration and scaling, WE simulations [28] were performed to enhance sampling of the ternary complex. WE operates by propagating a set of independent trajectories, referred to as walkers, each assigned a statistical weight. At regular intervals, walkers in under-sampled regions are cloned, while those in over-sampled regions are merged, ensuring efficient exploration of conformational space without biasing the dynamics [53]. In conventional WE simulations, the cloning and merging decisions (collectively referred to as “resampling”) are often based on binning along a predefined collective variable (CV) that tracks the progress toward a transition of interest. However, in some systems, a single CV is insufficient to capture the processes of interest. To overcome this challenge, we utilized an alternative algorithm for resampling known as REVO [18]. REVO is a bin-less method that prioritizes trajectories using an objective quantity called the “trajectory variation”, which is typically defined as sum of all-to-all distances among the walkers [18]. In our case, the distance metric is defined as

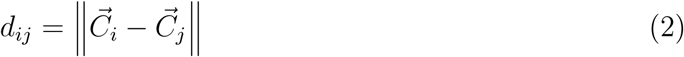

where 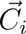 represents a vector of distances between a common set of atom pairs as measured in walker i. At each resampling interval, the variation between walkers is calculated according to

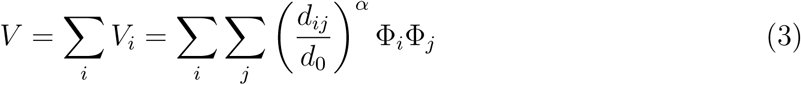

where V*_i_* r epresents a variation associated with the walker i, d*_ij_* denotes the distance between walker i and j as determined by the distance metric, and α is the exponent that balances the relative strength of the distance and the novelty functions; here, α = 4. Φ is the novelty function, which quantifies the relative importance of each walker, and here is defined as Φ*_i_* = C log w*_i_*/ log p_min_ where p_min_ = 10*^−^*^12^ and C = 100 following previous work [15, 25, 16]. The term d_0_ serves to make the variation function unit-less and to allow comparison between different distance metrics, but does not affect resampling behavior.

REVO works iteratively in each resampling phase: it first proposes a pair of walkers to merge and a single walker to clone. The expectation value of V is calculated both before and after this coupled cloning and merging step. If V is expected to increase, the move is carried out and another move is proposed. This continues until the variation is no longer expected to increase. Importantly, the surviving walker for each cloning operation is only chosen after a given move is accepted. A value of p_max_ = 0.1 was employed that prevented merging between pairs of walkers whose combined weight would exceed p_max_. WE simulations were carried out in the isothermal–isobaric (NPT) ensemble at 310 K with a timestep of 4 fs.

We utilized 31 atom pairs at the DCAF15-RBM39 interface to define the distance metric (Table S1). A larger set of pairs were first defined using all Cα–Cα residue pairs within 8 Å in the equilibrated apo crystal structure. The apo structure was then simulated for 500 cycles using WE with 48 walkers and 2500 steps (10 ps) per cycle. From these trajectories, we calculated the range of pairwise distances sampled and those that exhibited large distances (Δd > 0.4 nm) were removed, leaving only the more stable, persistent interface contacts. WE simulations using this reduced set of contacts were found to consistently generate unbinding trajectories within 500 cycles.

## Results

To determine an appropriate scaling factor λ for modifying inter-protein interactions, we performed WE simulation using the apo system and ran with four different simulation setups, λ = 0.00, 0.05, 0.10, and 0.15. Each simulation was run for 500 cycles using 48 walkers and 2500 steps per cycle and was replicated 3 times (with the exception of λ = 0.15, which was replicated 10 times) to assess reproducibility of the results. The weighted average of the pairwise distances between aforementioned atom pairs were calculated and analyzed as a probability distribution of distances. Figure 2 represents the distances between the two proteins in our system along with their likelihood. At λ = 0, corresponding to full interaction strength, the complex remained tightly bound throughout the simulations, with minimal sampling of dissociation. As λ increases and the interprotein interactions are reduced we observe a gradual broadening of the probability distribution towards higher distances. At the highest scaling level we examined, λ = 0.15, we observed dissociation of the protein complex. This scaling factor was therefore selected for all subsequent simulations.

**Figure 2:**
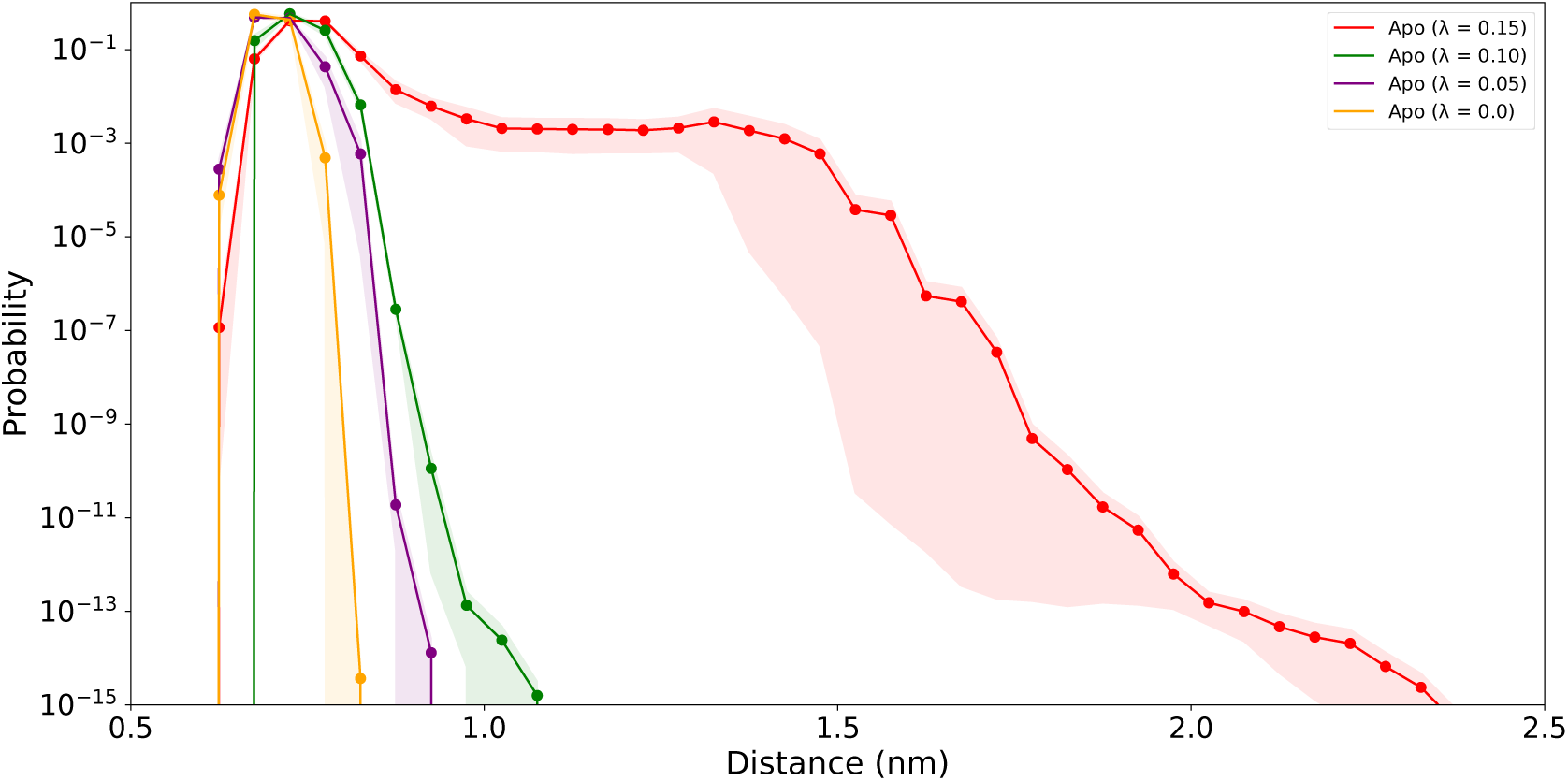
Probability distribution for the Apo form of DCAF15–RBM39 system under varying λ values. Shaded regions represent the standard error of the mean. Increasing λ enhances sampling of dissociated states up to λ = 0.15.

WE simulations for the molecular glue systems were replicated 10 times, and subjected to the same analysis as the scaling factor optimization simulations with Apo system (Figure 3). Using E7820 and the Apo system as benchmarks for high-potency and no-glue control conditions, respectively, the performance of the other glue molecules followed expected trends in accordance with their EC_50_ values. Low-potency glues exhibited a greater likelihood of dissociation, whereas high-potency glues were more likely to maintain a stable ternary complex.

**Figure 3:**
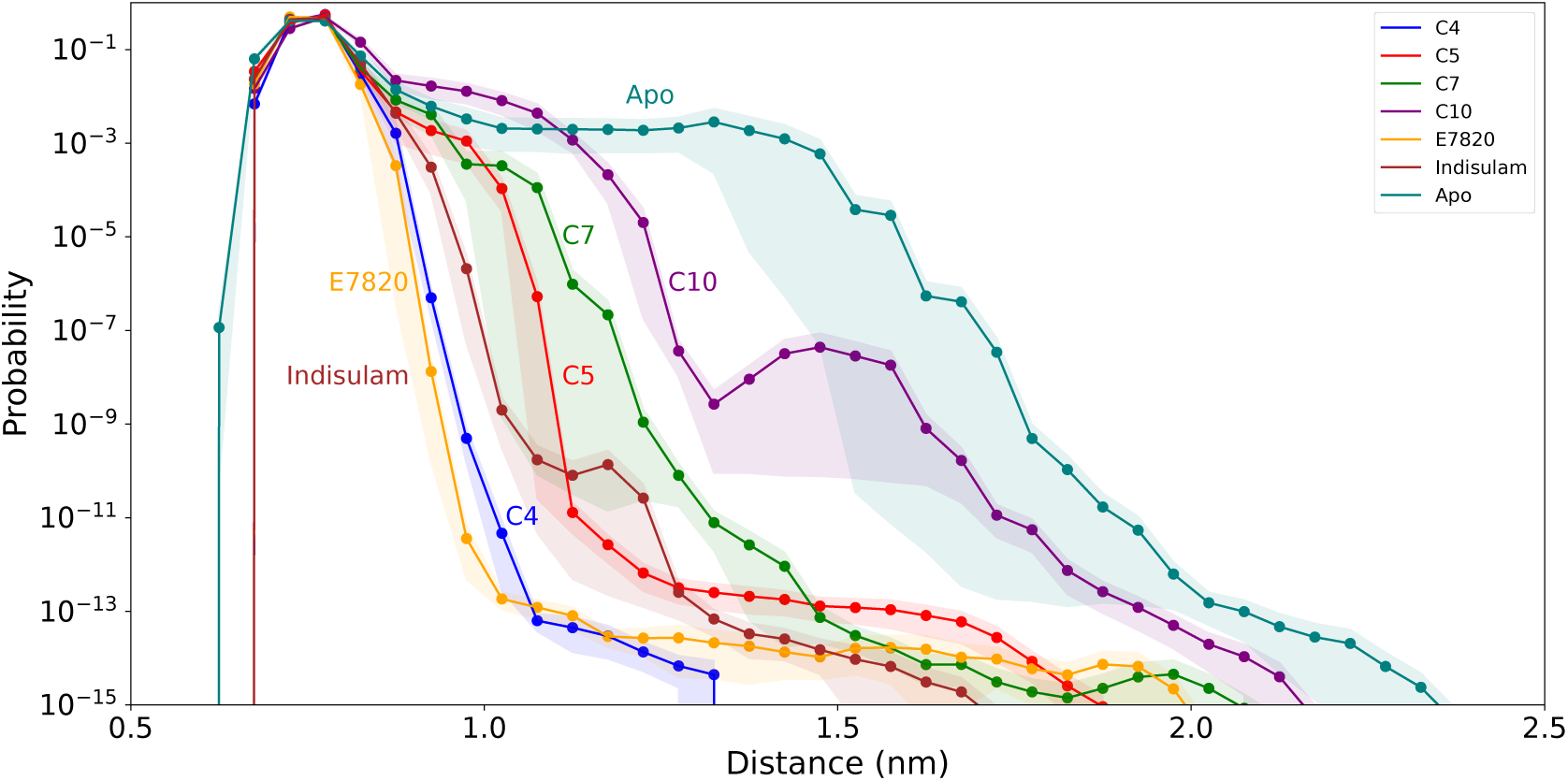
Probability distributions of the average interface distances for DCAF15–RBM39. Interface distances are calculated using the atom pairs mentioned previously. Probabilities are computed from the trajectory weights, normalized, and plotted on a logarithmic scale. Shaded regions represent the standard error of the mean.

To evaluate the molecular glue performance, WE simulations of different glue systems were replicated 10 times and subjected to the same analysis as the scaling factor optimization simulations with Apo system. Unbinding was quantified using the average interprotein distance (d_mean_) computed over the same set of residues that was used for distance metric calculations. We observed that E7820, C4 and Indisulam were less likely to explore farther distances, with Insidulam having larger uncertainty below probabilities of 10*^−^*^9^ (Figure 3). The C5 and C7 ligands were observed to sample larger distances, around d_mean_ = 1.2 nm, with a greater probability. C7 trajectories exhibited large variation in this region, making it difficult to rank C5 and C7 accurately among each other. Lastly, the non-degrader C10 clearly showed the highest likelihood to explore farther distances despite the high uncertainty in probability for distances between 1.3 and 1.7 nm.

To summarize this data, Figure 4 shows the average interface distances at a fixed probability threshold of 10*^−^*^8^ in comparison to their previously published EC_50_ values. This probability threshold value was chosen to represent the region of the distribution where systems exhibit clear separation and where probability estimates are stable across independent WE simulations. In the literature, C5 has an EC_50_ of > 50 µM, while C10 shows no measurable potency. In Figure 7, these compounds were plotted at 50 µM and 100 µM, respectively. This was done to reflect their relative activity while allowing comparison with other glues. This comparison shows that we are able to correctly rank order most of the glue molecules, with the exception of C5 and C7. A linear fit shows a Pearson correlation coefficient of R^2^ = 0.92.

**Figure 4:**
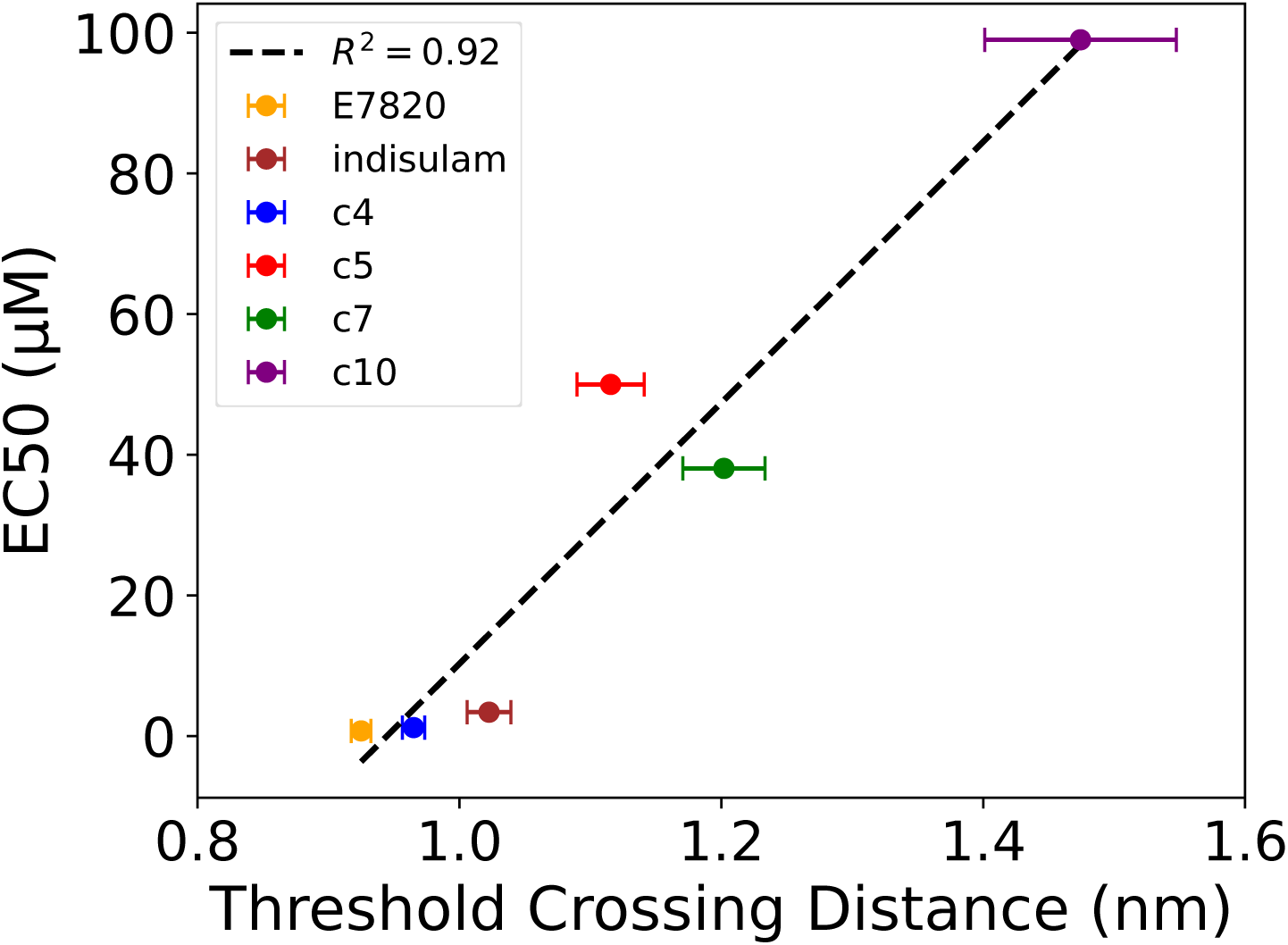
Relationship between experimental EC_50_ values and the DCAF15-RBM39 interface distance at which the probability distribution (Figure 3) crosses 10*^−^*^8^ threshold. Error bars represent the standard error of the mean of the threshold-crossing distances computed across independent WE simulations. EC_50_ values for C5 and C10 were plotted at 50 µM and 100 µM, respectively, to reflect low or unmeasurable potency while enabling comparison with other molecular glues. A linear fit yielded R^2^ = 0.92.

To understand the differences between the unbinding of degrader and non-degrader glue molecules, we have compared the bound and unbound poses of E7820 and C10 systems (Figure 5). For structures at high distances, we chose the highest probability walkers with d_mean_ ≥ 1.5 nm. As expected, C10 had a higher likelihood of reaching the unbound state compared to E7820, and in both trajectories, the glue molecules showed a preference towards DCAF15 after complex dissociation. To further examine the persistence of interactions between the glue molecules and the proteins, we computed the frequency of the interactions across all WE runs (Figure 6 and Table S2). Interactions were defined as protein residues within 0.25 nm of the glue molecules, and we focused on interactions that occur with a probability of at least 0.1, after taking the WE weights of the observed structures into account. We observed the indole ring of the glue molecules contributes the most to proteinglue interactions. In all systems, Asp264 on RBM39 was able to establish hydrogen bonds with indole and sulfonamide nitrogens and were the most dominant interactions, however we observed at least a 3-fold drop in frequency of these interactions in C10 compared to E7820 (Table S2). Some other notable interactions were the hydrogen bond between the indole nitrile (for E7820 and C4) and Arg552 sidechain hydrogens, which was replaced with Chlorine (for Indisulam and C10) and resulted in a 5 fold increase in interaction frequency along with the addition of Ile269 proximity. Substituting the methyl group of the indole ring (for E7820) with a hydrogen dropped the Val556 contact frequency by up to half for C10. Also, as C10 lacks of central sulfonamide, Pro233 proximity was less frequent and was below the 0.1 frequency threshold.

**Figure 5:**
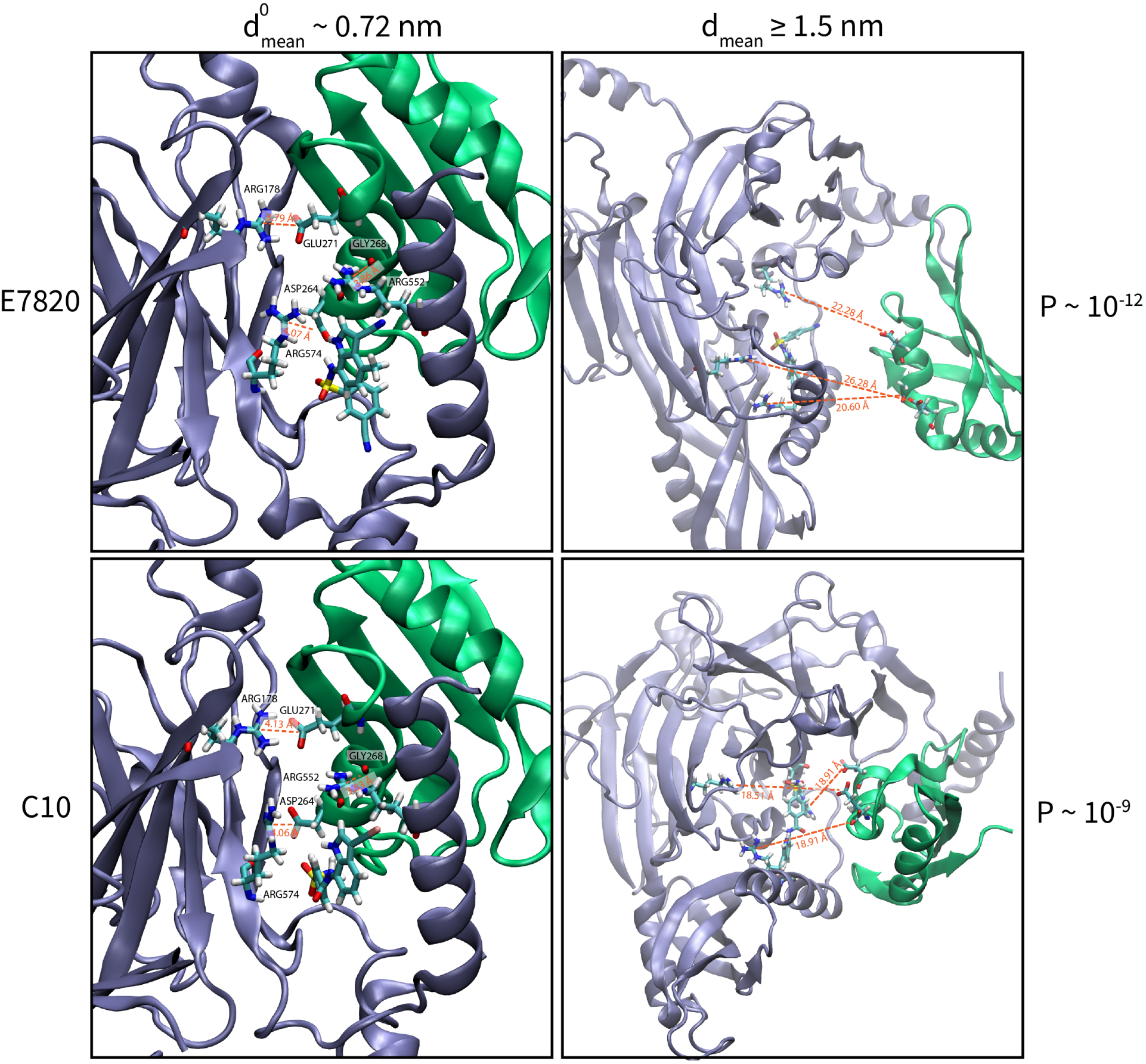
Snapshots from WE simulations shown in VMD [29]. E7820-bound and C10-bound ternary complex showing the initial pose from the side view (left) and the final pose from the top view (right) of the walker with the highest statistical weight after crossing the 1.5 nm threshold in average interface distance. DCAF15 is shown in purple, RBM39 in green, and glue molecules in licorice representation. Selected interface residues are labeled (left) and shown in licorice representation. Distances between a subset of interacting pairs are indicated with orange dashed lines.

**Figure 6:**
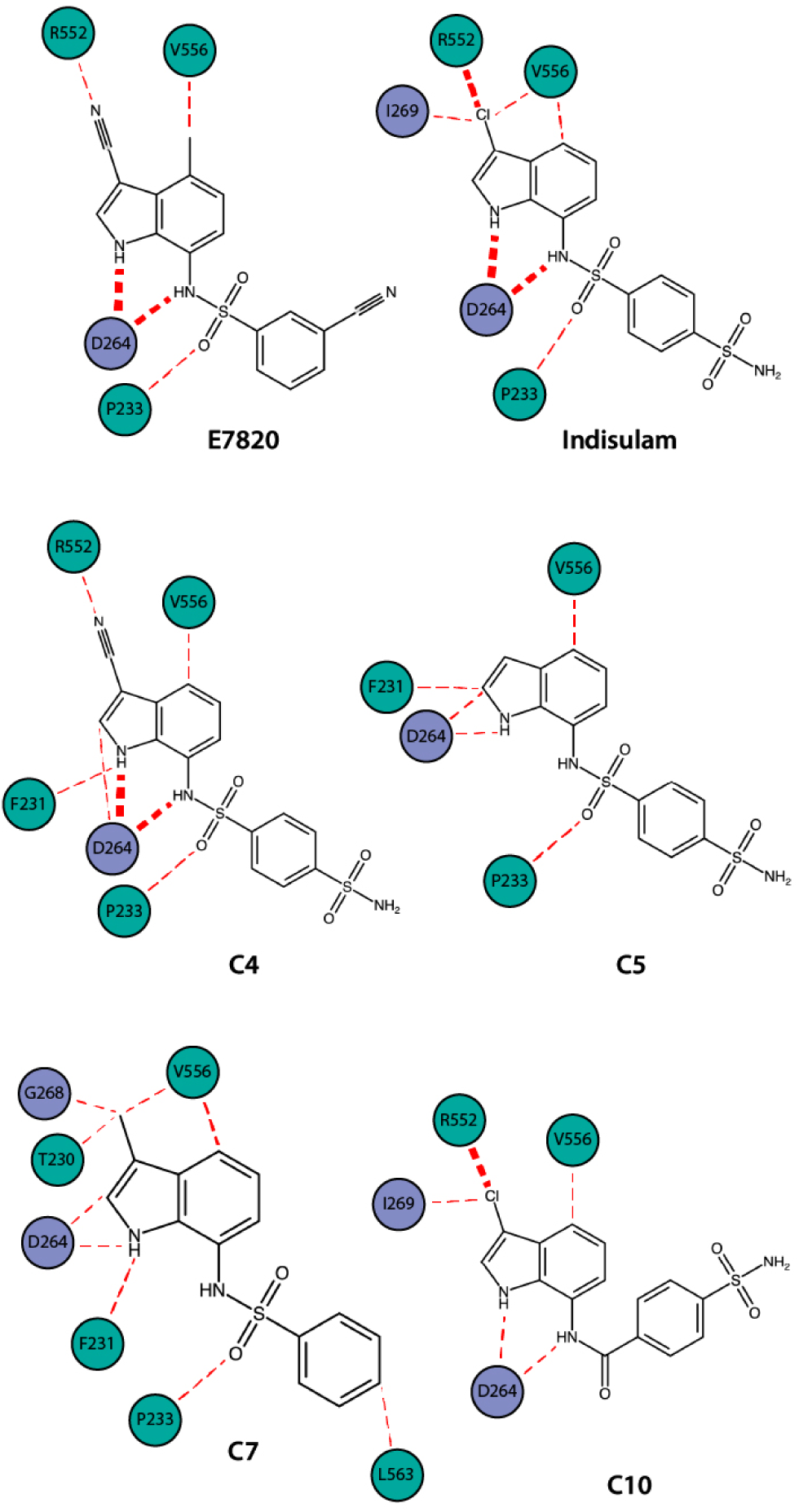
Comparison of interaction frequencies of E7820, Indisulam, C4, C5, C7, and C10 across all WE simulations. DCAF15 and RBM39 residues are shown in green and blue, respectively. Interaction frequencies are represented by the thickness of red dotted lines, with thicker lines indicating higher frequencies; line length is arbitrary and does not provide additional information.

We next perform a residue-level contact analysis to reveal the stability of important interactions between DCAF15 and RBM39 (Figure 7). These pairs include known salt bridges and hydrogen bonds reported by Bussiere et al.[10], supplemented with additional contacts identified by visual inspection of the simulation trajectories. From these pairs, we selected the carbon atoms nearest to the interacting atom pairs to allow some freedom of movement (Table S3). To improve the visualization, we estimated the interatomic distances between the selected carbon atom pairs at frames in which the corresponding interactions were formed (d_eq_). We then subtracted d_eq_ from the computed distances to plot ΔDistance in the figure. The negative log probability of these distances are computed across all WE simulations for a given system. Two interaction pairs that are within proximity of the glue indole ring stood out: Arg552(DCAF15)-Gly268(RBM39) and Arg574-Asp264. Notably, the Arg552-Gly268 and Asp557-Ala317 interactions are maintained even in the absence of a glue. The Arg574-Asp264 interaction is moderately stable in the apo system. Structural studies by Faust et al. have shown that this interaction forms as a salt-bridge in the absence of the glue molecule and is involved in a water-mediated hydrogen-bond in the presence of E7820, corresponding to our observed ΔDistance of 0.3 in the E7820 system. This interaction plays a crucial role in anchoring the ternary complex and is thought to contribute to the overall pharmacophore. The C4 systems also maintained closer Arg574-Asp264 distances, while weaker glues such as C10, which lacks the central sulfonamide, showed a significant destabilization in this interaction. The absence of this hydrogen bonding capability in C10 likely contributes to the observed lack of ternary complex stability and aligns with its lack of biological activity. In addition to this, a strong interaction between Arg552 and Gly268 were observed in our simulations. Previous mutational studies demonstrate that substitutions at Gly268(e.g., G268V or G268W) prevents the RBM39 recruitment and the formation of a ternary system. This suggests that Gly268 plays a critical role in stabilizing the DCAF15RBM39 interface. The simulation data in this study further supports this observation: in systems with active glue molecules (e.g., E7820 and C4), the Arg552-Gly268 contact remains stable throughout the trajectory. In contrast, in the inactive glue system of C10, this interaction becomes less likely or fully disrupted. Particularly for Indisulam, we observe significant stabilization of Met169-Arg275 and Met170-Arg275 contacts in comparison to apo. While we did not observe direct stabilizing interactions between the glue molecule and these residues, it is possible that indisulam allosterically stabilizes these hydrogen bonds.

**Figure 7:**
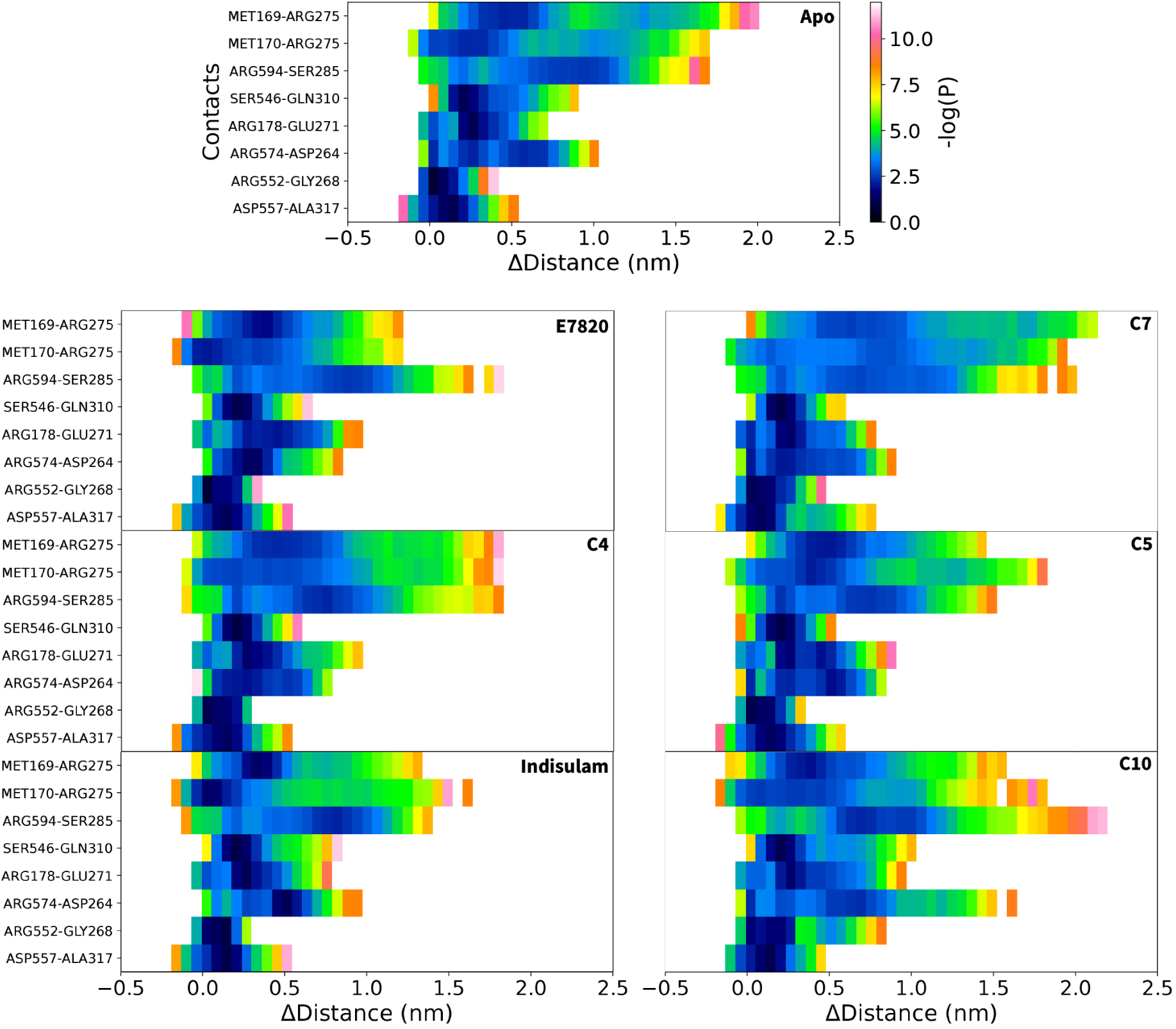
Heatmaps of DCAF15-RBM39 contact-distance distributions for all glue systems. Each row corresponds to a contact and the x-axis represents the change in distance relative to the bound pose. Columns are colored according to the probability of these distance changes: darker regions (blue) indicate high-probability states, whereas lighter regions (white) indicate low-probability or unobserved distances.

## Discussion and Conclusion

In this study, we developed a computational workflow to evaluate molecular glue candidates and captured their relative performance in a manner that aligned with experimentally measured EC_50_ values. Ternary complexes formed with high-potency glues, such as E7820 and C4, were observed to be more likely to show stable contacts while weaker or inactive glues, including C10, exhibited increased probabilities of dissociation and weaker interactions along key hydrogen bonds. We were able to correctly rank-order high potency glues such as E7820, C4 and Indisulam, and correctly identify the least stable complexes such as the Apo system and the non-degrader C10 glue molecule. This is encouraging for the ability of the method to assess the performance of high potency glue molecules. However, for glues with intermediate potencies like C5 (EC_50_ > 50 µM) and C7 (EC_50_ = 38.06 µM), the method shows a greater variation in the probability distributions, leading to noisier predictions of relative efficacy. Additional data will be required to assess the reliability of rank ordering in this mid-range activity window.

To generate trajectories of complex unbinding, we utilized the WE enhanced sampling method. Other enhanced sampling algorithms, such as those that introduce external biasing potentials, could also be used for this purpose. Although the comparison of raw computational cost between WE and bias potential-based methods can be challenging, WE can be more straightforward to apply in practical terms. For example, unlike umbrella sampling (US) [48], which requires a careful selection of umbrella windows, force constant, and initial configurations for each window, our WE approach (REVO [18]) only requires the definition of a distance metric between walkers and a single initial conformation of the bound complex. We note, however, that conceptually similar progressive initialization schemes can also be implemented within an Umbrella Sampling framework [14]. The most significant difference between the two approaches is in the computation of the probability distribution. In WE, the distributions are directly calculated from the statistical weights associated with the walkers. Weights of the low-probability states are a function of entire trajectory histories, taking into account how many times a given trajectory has been replicated. In contrast, methods like US construct probability distributions by stitching together biased trajectory ensembles and making assumptions about the overlap which are not always satisfied. For instance, even if trajectories from adjacent sampling windows show overlap in the biasing variable, they might still be separated along an orthogonal barrier that is not captured by the biasing variable. Without a continuous trajectory that spans from the bound state to the unbound state, there is no guarantee that all orthogonal barriers have been crossed [59].

This study can be extended to new systems by following the same simulation protocol and parameter values, but a set of validation simulations with the Apo complex – and a strong glue molecule complex if available – should be performed to assess sampling of the unbound state and the variations between runs. Here we relied on a common scaffold to determine suitable starting poses for each compound inside the binding pocket, which allowed us to assess the glues in a ternary complex without the need for separate experimental structures for each glue molecule. This resulted in the glue molecule being the only variable that distinguishes the systems. We note that the Indisulam binding pose we generated from heavy atom overlap with the E7820 pose differed only slightly from the Indisulam crystal structure (PDBID: 6SJ7) with a heavy-atom RMSD of 0.8647 ^°^A after alignment to the binding pocket (Figure S1). Although small errors in the starting poses could be naturally corrected over the course of the WE simulations, larger errors would likely result in an under-estimation of a compound’s stabilizing effect.

The simulation framework employed here enabled effective exploration of ternary complex stability between DCAF15 and RBM39 with different glue molecules. Average and pair-based contact analysis replicated important interactions reported in previous studies, such as Arg552-Gly268 and Arg574-Asp264, that were stabilized or disrupted depending on the glue molecule. We believe these findings support the utility of enhanced sampling simulations as an effective computational approach for evaluating ternary complex stability in the presence of molecular glues. This framework offers a quick and straightforward method for assessing glue candidates and provides a foundation for future efforts in molecular glue design.

## Data and Code Availability

Python scripts used to scale PPI interactions and run WE simulations, as well as the Conda environment YAML file are available at https://github.com/berkatik/glue.git.

