## Supplemental Info for "Assessing the Stability of Molecular Glues with Weighted Ensemble Simulations"

Table S1: Residue-level interface contacts between RBM39 and DCAF15 used for the distance metric. All the distance calculations are done between $C_\alpha$ atoms.

| RBM39 | DCAF15 |
|---|---|
| Asp264 | Thr230 |
| Asp264 | Phe231 |
| Asp264 | Gln232 |
| Met265 | Thr230 |
| Arg267 | Thr230 |
| Gly268 | Pro229 |
| Gly268 | Thr230 |
| Gly268 | Phe231 |
| Gly268 | Ser549 |
| Gly268 | Arg552 |
| Ile269 | Trp548 |
| Ile269 | Ser549 |
| Ile269 | Ser550 |
| Ile269 | Arg552 |
| Ile269 | Lys553 |
| Phe270 | Ser549 |
| Glu271 | Phe228 |
| Glu271 | Ser549 |
| Pro272 | Tyr226 |
| Pro272 | Pro227 |
| Pro272 | Phe228 |
| Pro272 | Ser544 |
| Pro272 | Gly545 |
| Pro272 | Ser546 |
| Pro272 | Trp548 |
| Pro272 | Ser549 |
| Phe273 | Gly545 |
| Leu316 | Lys553 |
| Leu316 | Ser554 |
| Leu316 | Val556 |
| Ala317 | Val556 |

Table S2: Interaction frequencies between glue molecules and proteins (Figure 6).

| Atom-Residue Pairs | Frequency |
|---|---|
| E7820:H5 – DCAF15:Val556 | 0.221 |
| E7820:H4 – DCAF15:Val556 | 0.221 |
| E7820:H3 – DCAF15:Val556 | 0.211 |
| E7820:O1 – DCAF15:Pro233 | 0.160 |
| E7820:N4 – DCAF15:ARG552 | 0.129 |
| E7820:H12 – RBM39:ASP264 | 0.739 |
| E7820:H8 – RBM39:ASP264 | 0.484 |
| Indisulam:Cl – DCAF15:ARG552 | 0.542 |
| Indisulam:O1 – DCAF15:PRO233 | 0.160 |
| Indisulam:H2 – DCAF15:VAL556 | 0.157 |
| Indisulam:Cl – DCAF15:VAL556 | 0.134 |
| Indisulam:H10 – DCAF15:ASP264 | 0.710 |
| Indisulam:H7 – DCAF15:ASP264 | 0.558 |
| Indisulam:Cl – DCAF15:ILE269 | 0.122 |
| C4:H2 – DCAF15:VAL556 | 0.172 |
| C4:O1 – DCAF15:PRO233 | 0.160 |
| C4:N3 – DCAF15:ARG552 | 0.151 |
| C4:H10 – DCAF15:PHE231 | 0.127 |
| C4:H10 – RBM39:ASP264 | 0.585 |
| C4:H7 – RBM39:ASP264 | 0.395 |
| C4:H4 – RBM39:ASP264 | 0.104 |
| C5:H3 – DCAF15:VAL556 | 0.206 |
| C5:O1 – DCAF15:PRO233 | 0.165 |
| C5:H11 – DCAF15:PHE231 | 0.128 |
| C5:H5 – RBM39:ASP264 | 0.191 |
| C5:H11 – RBM39:ASP264 | 0.116 |
| C7:H2 – DCAF15:VAL556 | 0.260 |
| C7:H11 – DCAF15:PHE231 | 0.183 |
| C7:O1 – DCAF15:PRO233 | 0.154 |
| C7:H13 – DCAF15:VAL556 | 0.130 |
| C7:H12 – DCAF15:VAL556 | 0.124 |
| C7:H14 – DCAF15:VAL556 | 0.123 |
| C7:H12 – DCAF!5:THR230 | 0.109 |
| C7:H10 – DCAF15:LEU563 | 0.108 |
| C7:H4 – RBM39:ASP264 | 0.124 |
| C7:H12 – RBM39:GLY268 | 0.118 |
| C7:H14 – RBM39:GLY268 | 0.118 |
| C7:H13 – RBM39:GLY268 | 0.113 |
| C7:H11 – RBM39:ASP264 | 0.109 |

Table S2 (continued): Interaction frequencies between glue molecules and proteins (Figure 6).

| Atom-Residue Pairs | Frequency |
|---|---|
| C10:Cl – DCAF15:ARG552 | 0.601 |
| C10:H2 – DCAF15:VAL556 | 0.113 |
| C10:H6 – RBM39:ASP264 | 0.211 |
| C10:H5 – RBM39:ASP264 | 0.156 |
| C10:Cl – RBM39:ILE269 | 0.132 |

Table S3: Residue-level interface contacts between RBM39 and DCAF15 used for distance calculations in Figure 7.

| RBM39 | DCAF15 |
|---|---|
| Glu271:CD | Arg178:CZ |
| Asp264:CG | Arg574:CZ |
| Ser285:C | Arg594:CZ |
| Ala317:CA | Asp557:CG |
| Arg275:CZ | Met170:CA |
| Arg275:CZ | Met169:CA |
| Gly268:C | Arg552:CZ |
| Gln310:CG | Ser546:CB |

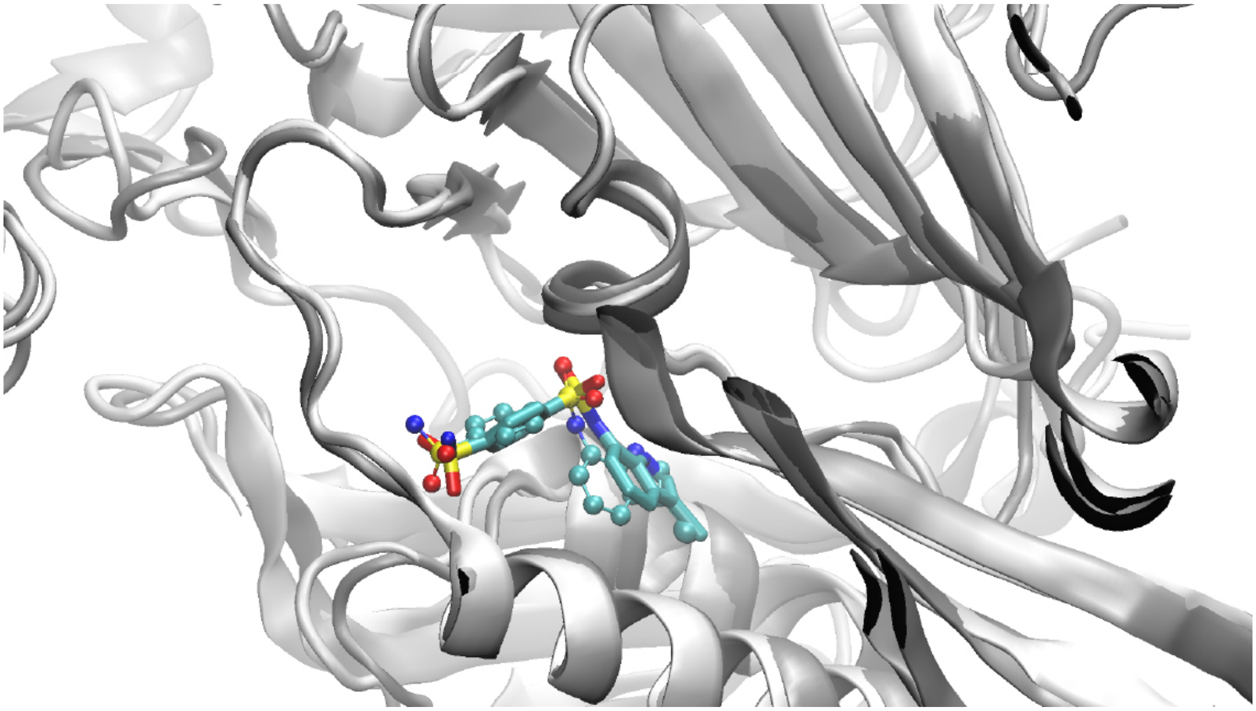

Figure S1: Comparison of the superimposed Indisulam crystal structure (PDB ID: 6SJ7), shown in CPK representation, and the structure generated using the CHARMM-GUI Ligand Reader & Modeler, shown in licorice representation (RMSD = 0.8647 Å ).
